# Functionally unique, specialised, and endangered (FUSE) species: towards integrated metrics for the conservation prioritisation toolbox

**DOI:** 10.1101/2020.05.09.084871

**Authors:** J. N. Griffin, F. Leprieur, D. Silvestro, J. S. Lefcheck, C. Albouy, D. B. Rasher, M. Davis, J.-C. Svenning, C. Pimiento

## Abstract

Identifying species with disproportionate contributions to biodiversity can lead to effective conservation prioritisation. Despite well-established methods for identifying endangered species adding inordinately to *evolutionary* diversity, in this context *functional* diversity has been overlooked. Here, we compare different metrics designed to identify threatened species that contribute strongly to functional diversity. We use the diverse and threatened global marine megafauna as a case study. We found that functional contributions of species are not fully captured in a single metric. Although we found a very strong correlation between functional specialisation and distinctiveness, functional uniqueness was only moderately correlated with the other two metrics and identified a different set of top-10 species. These functional contributions were then integrated and combined with extinction risk to identify species that are both important contributors to functional diversity *and* endangered. For instance, the top-10 *F*unctionally *U*nique *S*pecialized and *E*ndangered (FUSE) species contains three critically endangered, five endangered and two vulnerable species which - despite comprising only 3% of species - are among the top 10% most functionally unique and hold 15% of the global functional richness. The FUSE index was remarkably robust to different mathematical formulations. Combining one or more facets of a species contribution to functional diversity with endangerment, such as with the FUSE index, adds to the toolbox for conservation prioritisation. Nevertheless, we discuss how these new tools must be handled with care alongside other metrics and information.

## Introduction

Earth’s biodiversity is in crisis. Over the last half-century populations of many invertebrates (e.g. terrestrial insects, van Klink et al. 2020) and vertebrates (e.g. birds, Rosenberg et al. 2019) have plummeted, while global extinction rates are at least several orders of magnitude above historical background levels (Barnosky et al. 2011; Ceballos et al. 2015; Pimm et al. 2014). Many groups are imperilled across the terrestrial, freshwater, and ocean realms. For example, around 13% of birds, 40% of amphibians, and 30% of sharks and rays are currently threatened with extinction (Díaz et al. 2020). Given limited public and political will - and ultimately funding - to conserve Earth’s biodiversity, efforts have been made to prioritise threatened species. Yet while progress has been made to incorporate species’ contributions to phylogenetic diversity when triaging conservation goals (i.e., protecting evolutionarily distinct lineages; Isaac et al. 2007; Mooers et al. 2008), efforts to integrate the *functional diversity* afforded by each species has lagged.

Functional diversity is a dimension of biodiversity that captures variation in the traits and ecological roles of species (Tilman 2001; Violle et al. 2007). Traits such as body size, mobility and feeding behaviour determine an organism’s capacity to process resources and how it experiences and interacts with its environment. High functional diversity within a community allows species to exploit diverse resources and more efficiently consume and transport energy and nutrients within and across ecosystems (Dee et al. 2016; Gagic et al. 2015; Lefcheck and Duffy 2015).

Quantitatively, functional diversity can be defined as the distribution of species in a functional trait space whose axes represent combinations of traits (Rosenfeld 2002; Mason et al. 2005; Villéger et al. 2008; Fig. 1A). Functional diversity is now widely recognized as a multi-faceted entity (Villéger et al. 2008; Mouillot et al. 2013) with (1) functional richness defined as the amount of functional trait space filled by all species in a species assemblage, and therefore being strongly influenced by extreme trait values; (2) functional evenness representing how species are regularly distributed in the functional trait space; and (3) functional divergence describing how far species are from the centre of the functional trait space. These three complementary facets, all together, describe the distribution of species within the functional trait space (Mason et al. 2005; Villéger et al. 2008).

**Figure 1.**
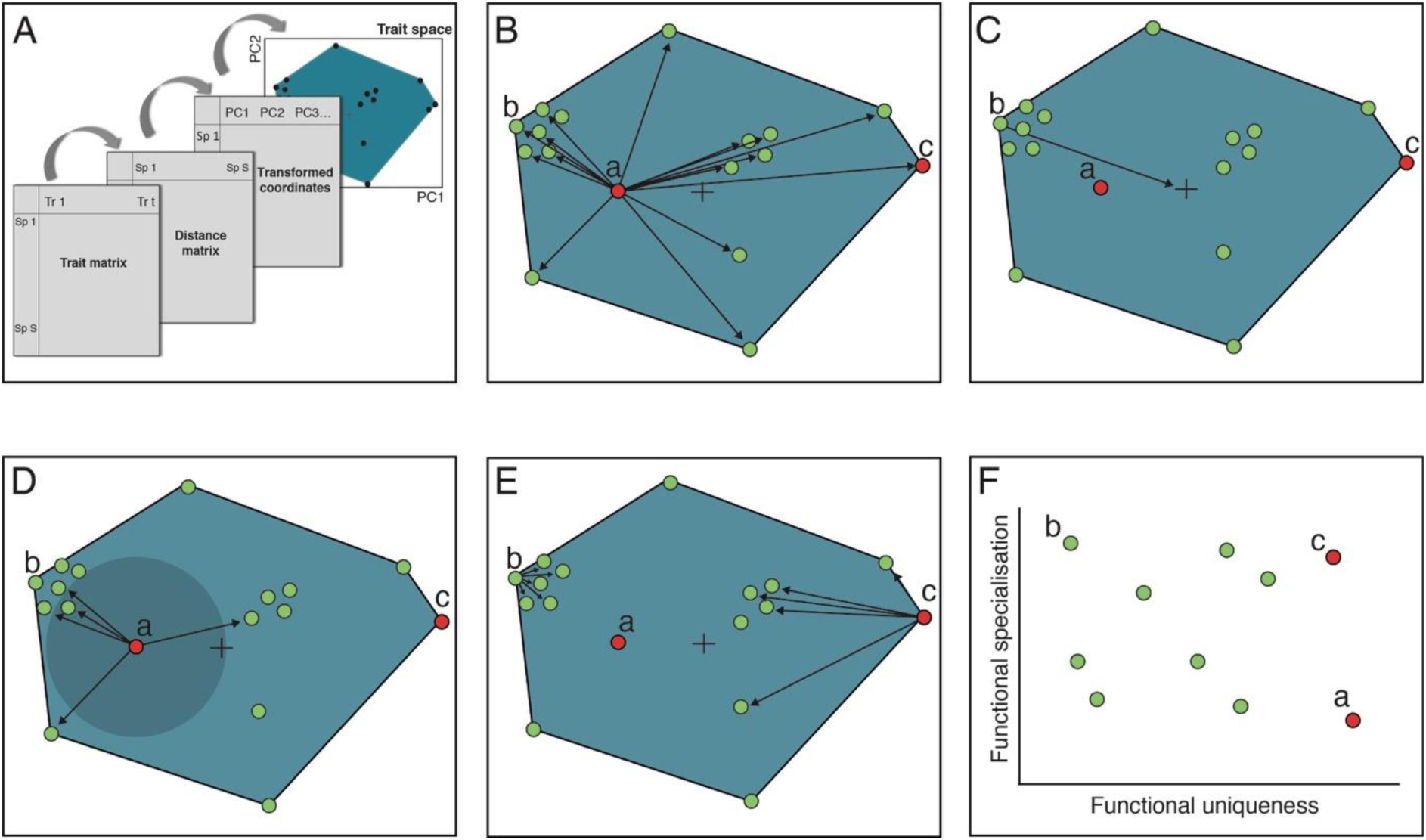
Species’ contributions to functional diversity and their endangerment. **A**) Following trait compilation, species x species trait distances are rescaled into a multidimensional ‘functional space’ (tables based on Villéger et al. 2008). **B**) Functional distinctiveness is the mean distance to all other species in the community, characterising a species’ *global* relative position in trait space, as shown for *species a*. **C**) Functional specialisation is measured as a species’ distance from the trait space centroid. Species with high functional specialisation (such as *species b*) support extreme trait values and combinations. **D**) Functional uniqueness is measured as the mean distance to a set of *n* nearest neighbours (here *n* = 5), characterising a species’ *local* relative position in trait space. Species with high functional uniqueness like *species a* are isolated in trait space so their loss would leave a large part of trait space unoccupied. **E**) Functionally unique and functionally specialised species are not necessarily the same: functionally unique species may have traits that are not extreme (*species a*), while functionally specialised species may have low functional uniqueness (*species b*). Highly functionally specialised *and* highly functionally unique species (*species c*) are located in extreme and sparsely occupied areas of trait space. **F**) Species’ functional contributions can be combined with conservation status (green = non-threatened, red = threatened; also in panels B-E) to identify species that are both highly unique and/or specialised as well as being endangered. In this example, *species b* is functionally specialised but not endangered, *species a* is functionally unique and endangered and *species c* is functionally unique, specialised, and endangered.

Similarly, species’ contributions to functional diversity can be described through a suite of metrics (Fig. 1; Mouillot et al. 2013). ‘Functional distinctiveness’ (i.e. the mean distance between a given species and all others in the studied pool of species, Fig. 1B) is commonly used (Violle et al. 2017; Cooke et al. 2020) to define the degree of functional differentiation between a species and all others. Other aspects of species’ positions may also determine their contribution to functional diversity. Species with extreme traits – such as a very large size – are ‘functionally specialised’ and strongly contribute to maintaining the limits of functional space (Leitao et al. 2016; Mouillot et al. 2013; Fig. 1C). Species with unusual – though not necessarily extreme – traits are ‘functionally unique’, and sit relatively isolated in trait space (Fig. 1D), lacking functionally similar species that could compensate for their functional role if they were to be lost (Bellwood et al. 2003; Leitao et al. 2016; Mouillot et al. 2013).

Where a species is both functionally specialised *and* functionally unique, they are positioned in extreme and isolated positions in trait space and may singularly support a substantial portion of trait diversity, analogous to a lone species on a long branch in a phylogenetic tree (Fig. 1E). The extent to which these alternative species-level metrics provide redundant versus complementary information on species’ contributions to functional diversity has not been well-studied (but see Mouillot et al. 2013 and Leitao et al. 2016 for their use in the context of species community responses to disturbances).

Just as all threatened species do not contribute equally to functional diversity; all species are not at equal risk of extinction. Therefore, prioritisation of species based on their contribution to functional diversity should simultaneously consider extinction risk (Fig. 1). The International Union for Conservation of Nature (IUCN) evaluate and categorize species’ risk based on their rarity, rate of population decline, population size, area of geographic distribution, and degree of population fragmentation. The IUCN status of each species can be interpreted as an ordered categorical variable denoting increasing threat (least concern, near threatened, vulnerable, endangered, critically endangered) or translated into a probability of extinction over a given time interval (Davis et al. 2018; Mooers et al. 2008).

Building on this framework, species’ extinction risk has been combined with evolutionary distinctiveness to identify ‘evolutionarily distinct and globally endangered’ (EDGE) species (Isaac et al. 2007). There have also been recent efforts to integrate information on species’ functional distinctiveness and rarity, as indicated by range size (Grenie et al. 2018; Kondratyeva et al. 2019; Violle et al. 2017), and to incorporate functional diversity into spatial conservation planning (Pollock et al. 2017). However, there have been limited efforts to combine functional contributions to the more encompassing assessments of risk provided by the IUCN (but see Hidasi-Neto et al. 2015). Doing so has the potential to identify species that are making pivotal contributions to functional diversity and at the same time are facing high levels of endangerment, although we note that this is not always true: some species may be at low risk of extinction but possess highly unusual traits, or be at high risk but possess traits shared by many other species. Furthermore, the two approaches based on the phylogenetic and functional components of biodiversity should be complementary as recent studies showed that functional distinctiveness and evolutionary distinctiveness were weakly related for birds, mammals (Cooke et al. 2020) and tropical reef fishes (Grenie et al. 2018). Thus, the combined information on extinction status and functional traits may help secure not only biodiversity but also the future of functioning ecosystems.

In a recent study, we explored future changes in functional richness of the world’s marine megafauna under different extinction scenarios (Pimiento et al. 2020). We also introduced the novel FUSE index (Functionally Unique, Specialized, and Endangered) that combines functional uniqueness, specialisation, and global endangerment to identify threatened species of particular importance for functional diversity. Marine megafauna (defined as organisms with a maximum reported biomass ≥45kg; Estes et al. 2016) consist of over 300 species globally, mainly comprising marine mammals, sharks and rays, and bony fishes. These species are well suited for investigating species’ functional contributions to nature, as, not only do they play a variety of important functional roles – from whales transporting nutrients to surface waters, to sea otters preventing in the collapse of coastal kelp forests – but one-third of these species are also threatened with extinction due to human exploitation or other activities (Estes et al. 2016; Pimiento et al. 2020).

Here, we further investigate ways to combine the functional dimension of biodiversity and the level of species’ endangerment using this earlier dataset of marine megafauna. Specifically, we aim to address the following questions: 1) To what extent do species-level metrics of functional contributions provide complementary vs. redundant information? 2) How sensitive are metrics combining functional contributions and extinction risk to alternative formulations? and 3) How well do functional metrics capture the collective contribution of high-scoring species to functional richness?

## Methods

### Species and traits

The 334 marine megafauna species were as provided by Estes et al. (2016) and used in Pimiento et al. (2020). Several of these species use the marine environment marginally (i.e., brackish waters). The following 10 functional traits were assigned to each species: maximum body mass, thermoregulation, terrestriality, habitat zone, vertical position, migration, feeding mechanism, diet, breeding site, and group size/sociality. These traits were selected to capture the various ecological roles of megafaunal species. Full details and justification of trait selection and treatment are provided in Pimiento et al. (2020). In addition to traits, IUCN Red List status were gathered from the IUCN website in April 2019.

There were 51 species with one or more missing traits, which we inferred using multiple imputations based on the frequency of the trait combinations in each taxonomic order. At the same time, the IUCN status of the 24 ‘not evaluated’ (NE) species and the 48 ‘data deficient’ (DD) species were also inferred by resampling each missing status based on the distribution of the other IUCN status in each order. Multiple imputations for traits and IUCN status were run simultaneously for 1,000 iterations (see Pimiento et al. 2020 for further details).

### Construction of the trait space

A species trait distance matrix was created using a modified version of Gower’s distance (‘dist.ktab’ function of the *ade4* package in the R statistical software; R Development Core Team 2017), which allows the treatment of various types of variables, i.e., continuous, ordinal, nominal, multichoice nominal and binary (Pavoine et al. 2009). Using this matrix, a multidimensional Euclidean space was constructed based on a Principal Coordinates Analysis (PCoA) (Maire et al. 2015). The PCoA axes were retrieved using the ‘dudi.pco’ function (*ade4* package). We selected the first 6 PCoA axes that adequately represent the initial distance matrix to build the functional space according to the methods proposed by Maire et al. (2015). These axes represented 71.15% of the total inertia.

Note that although dendrogram-based methods have been historically popular in functional diversity studies (Petchey and Gaston 2002) and have been used to estimate individual species’ contributions to functional diversity (e.g. Hidasi-Neto et al. 2015, Pollock et al. 2017; and reviewed in Kondratyeva et al 2019), here we decided to use a multidimensional trait space approach that more effectively captures the underlying species’ functional distances (Maire et al. 2015). According to the mSD (mean squared deviation) metric proposed by Maire et al. (2015) a UPGMA dendrogram provided the worst representation of the initial Gower’s distances (see Fig S1), while a 6D trait space provides the most parsimonious representation (i.e. the best compromise between the quality of the representation of the initial distances and the number of axes, see Maire et al. 2015 for full methodological details and Pimiento et al. 2020 for application to the marine megafauna trait data).

### Resulting trait space

The trait space is described in detail in Pimiento et al. (2020) so we only briefly summarise it here to provide context. The first axis is strongly related to terrestriality (i.e., the ability to move between marine and terrestrial or riverine realms) and vertical position in the water column. The second axis of the trait space revealed a strong compartmentalization, with mammals occupying an area separate from that of fishes and sharks. The third axis of the trait space is mostly related to feeding mode, while the remaining three axes of the functional space are weakly associated with individual species traits and together represented 21% of the total inertia.

### Metrics of species contributions to functional diversity

Table 1 defines and outlines all the metrics used to characterize species’ contributions to functional diversity and its combination with level of global endangerment. For all the individual functional metrics (functional uniqueness [FUn], functional specialisation [FSp] and functional distinctiveness [Fdist]) the mean value across all imputations was used (see above and Pimiento et al. 2020). For each species, FUn was computed as the mean distance to the nearest neighbours, considering the one, three or five neighbours to test the sensitivity to the inclusion of varying portions of the trait-space neighbourhood (Mouillot et al. 2013; McWilliam et al. 2018). We focus on the case with 5 neighbours for consistency with Pimiento et al. (2020) but report results for all three cases in the supplementary materials to evaluate sensitivity. Species’ FSp was computed as the Euclidean distance of each species to the barycentre of the multidimensional trait space (Mouillot et al. 2013). We did not investigate the functional distinctiveness (Fdist) in Pimiento et al. (2020), but we include here as this metric is commonly used in other studies and is a general descriptor of a species’ mean dissimilarity to all others in the community (Grenie et al. 2018; Kondratyeva et al. 2019; Violle et al. 2017). Note that computing FUn and Fdist does not require the Euclidean trait space; rather, they can be directly calculated based on the Gower’s distance matrix (Kondratyeva et al. 2019). On the other hand, FSp does rely on the Euclidean trait space (i.e., the PCoA). In Pimiento et al. (2020), FSp was based on the 6-dimensional space; here we calculate FSp based on all the axes of the PCoA, which more faithfully reflects the original functional distances. Functionally uniqueness and specialisation (FUS) was calculated as the average of FUn and FSp, with the intention of identifying species with high values of both metrics. Averages can be dominated by individual components so may not indicate simultaneously high values across component parts (Byrnes et al. 2014). Therefore, we separately identified those species that are in the top 10% (90^th^ percentile) for both FUn and FSp (hereafter FUSq), before ranking these according to the mean value.

**Table 1.** Description of metrics used in this paper. References are: 1. Mouillot et al. (2013). 2. Violle et al. (2017). 3. Kondratyeva et al. (2019). 4. IUCN (2019). 5. Pimiento et al. (2020), 6. Mooers et al. (2008). Subscript P in acronyms refers to the alternative extinction probability-weighted version of the respective metric.

| Metric name | Acronym | Ref. | Calculation | What it measures |
| --- | --- | --- | --- | --- |
| Functional uniqueness | FUn | 1,2,3 | see *Methods* | Mean distance to set of neighbours; scaled 0-1 relative to others in the community |
| Functional specialisation | FSp | 1 | see *Methods* | Relative distance to functional space centroid; scaled 0-1 relative to others in the community |
| Functional distinctiveness | Fdist | 2,3 | see *Methods* | Mean distance to all other species in the community; scaled 0-1 relative to others in the community |
| Global endangerment | GE<sub>(P)</sub> | 4 | see refs. 4- 6 | Rank or probability of extinction risk based on the IUCN Red List; rank scaled 0-1; probability as in ref 6. |
| Functionally unique and specialised | FUS | This paper | (FUn + FSp)/2 | Species average uniqueness and specialisation; scaled 0-1 relative to others in the community |
| Top functionally unique and specialised | FUSq | This paper | see *Methods* | Species in 90th percentile for functional uniqueness and specialisation |
| Functionally unique and globally endangered | FUGE<sub>(P)</sub> | 5 | $\text{Ln}(1 + \text{FUn} * \text{GE}_{(P)})$ | Species functional uniqueness weighted by global endangerment; max. is $\text{Ln}(2)$ |
| Functionally specialised and globally endangered | FSGE<sub>(P)</sub> | 5 | $\text{Ln}(1 + \text{FSp} * \text{GE}_{(P)})$ | Species functional specialisation weighted by global endangerment; max. is $\text{Ln}(2)$ |
| Functionally distinctive and globally endangered | FDGE<sub>(P)</sub> | 5 | $\text{Ln}(1 + \text{Fdist} * \text{GE}_{(P)})$ | Species functional distinctiveness weighted by global endangerment; max. is $\text{Ln}(2)$ |
| Functionally unique, specialised and endangered | FUSE<sub>(P)</sub> | 5 | $\text{FUGE}_{(P)} + \text{FSGE}_{(P)}$ | Species with both high FUGE and high FSGE scores; max. is $\text{Ln}(4)$ |
| Functionally unique, specialised and endangered (alternative) | FUSE'<sub>(P)</sub> | This paper | $\text{Ln}(1 + \text{FUS} * \text{GE}_{(P)})$ | Species functional uniqueness and specialisation weighted by global endangerment; max. is $\text{Ln}(2)$ |
| Functionally unique, specialised and endangered (alternative B) | FUSE''<sub>(P)</sub> | This paper | $\text{Ln}(1 + (\text{FU} + \text{FSp}) * \text{GE}_{(P)})$ | Species functional uniqueness and specialisation weighted by global endangerment; max. is $\text{Ln}(3)$ |

All functional metrics (i.e., FUn, FSp, Fdist, FUS) and IUCN status (GE: LC=0, NT=1, VU=2, EN=3, CR=4) were scaled between zero and one to ensure equal contributions to products when combined with global endangerment in the metrics ‘Functionally Specialised and Globally Endangered’ (FSGE), ‘Functionally Distinctive and Globally Endangered’ (FDGE), ‘Functionally Unique and Globally Endangered’ (FUGE), and ‘Functionally Unique, Specialised and Endangered’ (FUSE). Note that the scaling of functional metrics and IUCN status was between zero and four in Pimiento et al. (2020), which affects the absolute, but not relative values of FUSE and related indices.

Metrics combining functional contributions and IUCN status (global endangerment; GE) included two alternatives: a) IUCN status treated as a relative score as in Pimiento et al. (2020), and here scaled between 0 and 1; and b) IUCN status treated as a probability of extinction over a 100 year time interval (Mooers et al. 2008). We acknowledge that there are recent advances in translating IUCN status to extinction probabilities (Andermann et al. 2020; Davis et al. 2018), but here use the probabilities from Mooers et al. (2008) as a first pass at the general sensitivity of our focal metrics to IUCN treatment. Weighting by relative score makes no assumptions about extinction risk probabilities and simply provides equal increments for each IUCN status. Weighting by probability of extinction on the other hand considers the non-linearity of risk with increasing IUCN status (Isaac et al. 2007, Mooers et al. 2008; Davis et al. 2018). Note that FUSE was originally proposed in Pimiento et al. (2020) as a simple sum of FUGE and FSGE and should not be interpreted in terms of a particular weighting of FUn, FSp and GE. Several alternative formulations of FUSE are possible: FUSE’, which can be interpreted as the mean of FUn and FSp (FUS) weighted by GE; and FUSE’’ which is the sum of FUn and FSp weighted by GE. As FUS, it is also desirable for FUSE to capture species that are both high in FUGE and FSGE. We repeated the process of identifying species in the top 10% of FUGE and FSGE, to identify FUSEq species.

## Analysis

To evaluate the correspondences between species’ rankings based on functional metrics we used bivariate scatterplots and Spearman’s rank correlation coefficient. After identifying and ranking the top ten highest scoring species for each metric, we summarised the overlap in species using the Jaccard similarity index. This simple metric is the ratio between the shared species to the total number of species across any given pairwise comparison.

To gauge the influence of species-level metrics on the global functional richness (the total volume of the 6D functional space as measured by the FRic index; Villéger et al. 2008), we removed the top-10 and top-20 highest-scoring species for each metric from the global pool of species and compared this to the removal of the same number of randomly selected species. The removal of 10 and 20 random species was each performed 1000 times.

## Results

### Redundancy and complementarity across metrics

Functional uniqueness correlated only moderately with functional specialisation and functional distinctiveness (Fig. 2) and identified a distinct set of top-10 species (Fig. S2). Note that these correlations weakened as fewer neighbours were considered in the calculation of functional uniqueness (Fig. S3). Functional specialisation (FSp) correlated very strongly with functional distinctiveness (Fdist), which explains why Fdist correlated strongly with FUS (Fig. S4), despite not being used directly in its calculation. Therefore, individual contributions are captured by functional uniqueness on the one hand, and functional specialisation *or* functional distinctiveness of the other hand.

**Figure 2.**
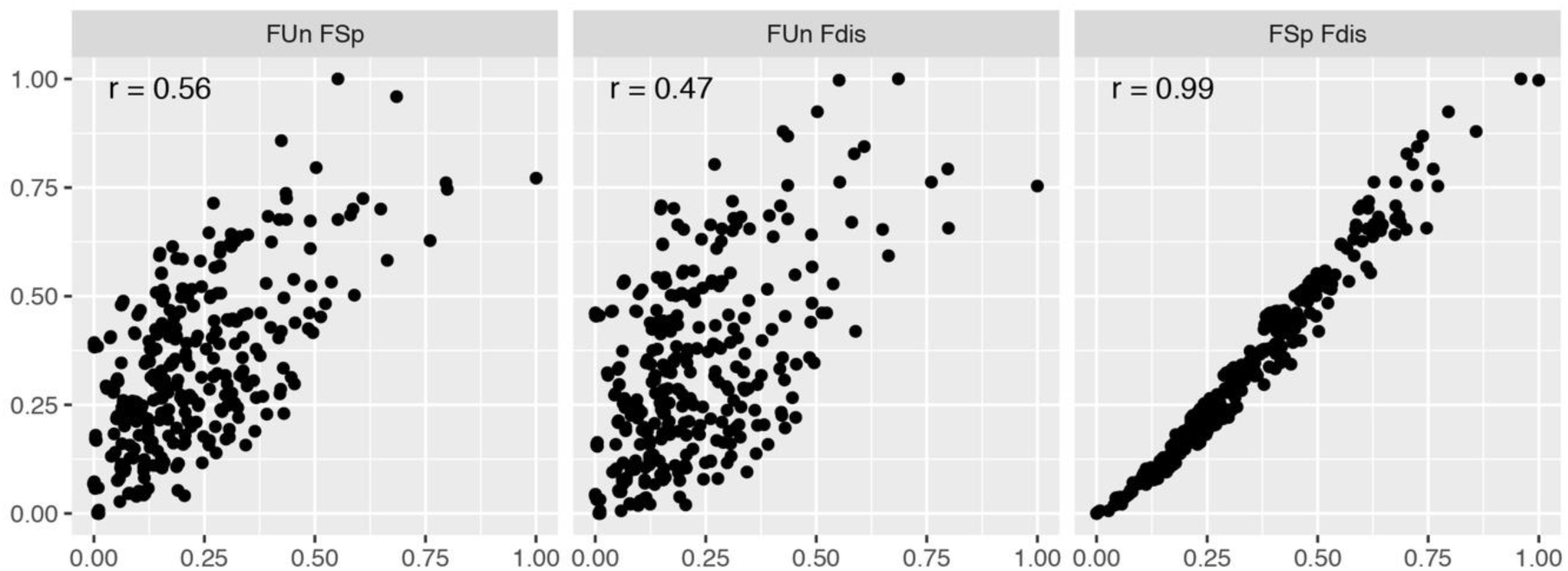
Correlations between metrics describing individual species’ contributions to functional diversity. FUn = functional uniqueness; FSp = functional specialisation; Fdist = functional distinctiveness. The Spearman’s rank correlation coefficient is denoted as r.

When integrating the functional indices weighted by IUCN status (with GE treated as a rank), the correlations between metrics were still apparent (0.90 ≤ r ≤ 1; Fig. S5). The top-ranked species under metrics including a weighting by IUCN status naturally contain a high proportion of threatened species (Fig. 3). For example, the top-10 FUSE species comprise three critically endangered, five endangered, and two vulnerable species. There is complementarity between those species identified using FUGE and FSGE, and less complementarity between FSGE and FDGE (Fig. 3; note panel E showing pairwise similarities between top-10 species sets).

**Figure 3.**
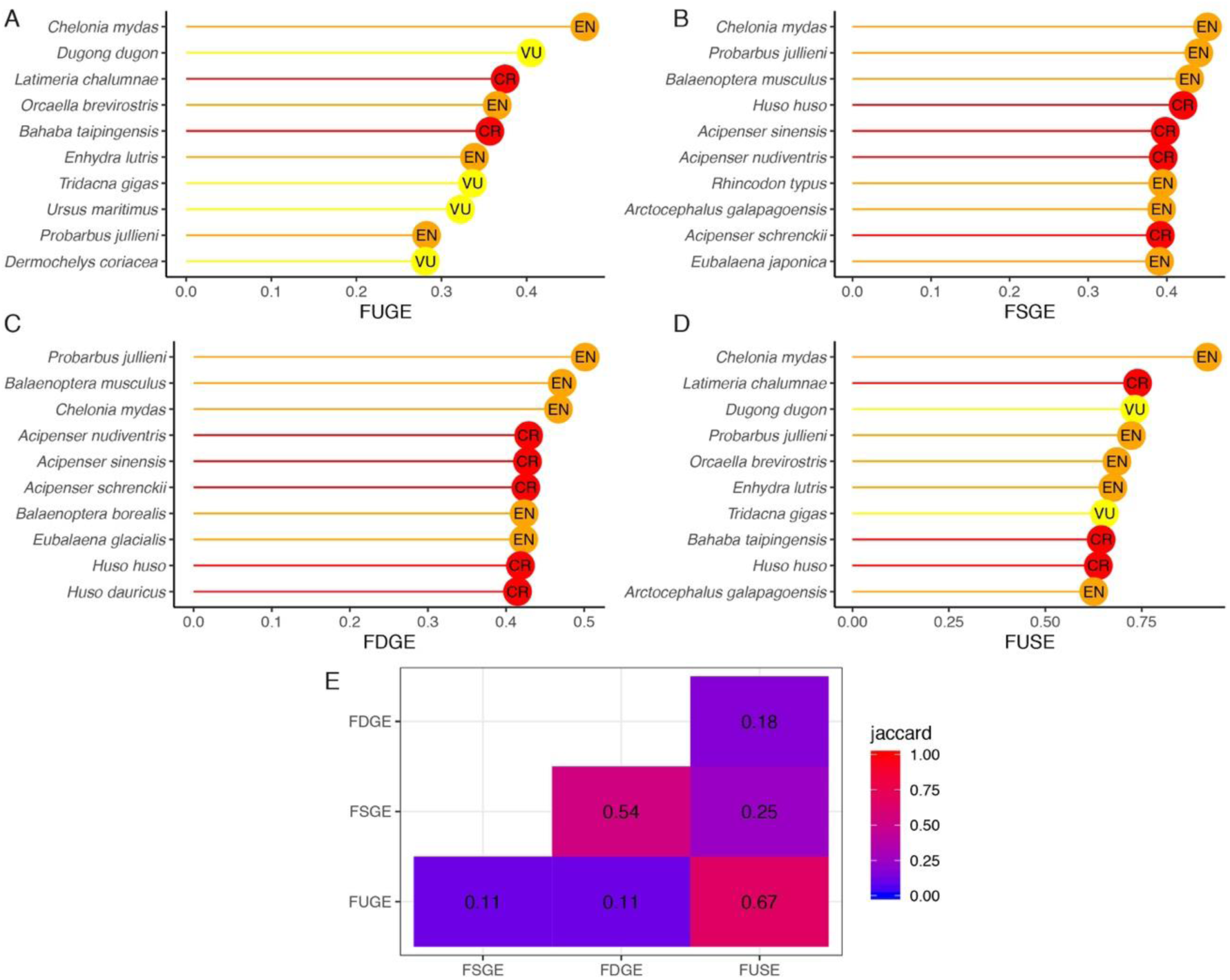
Top-10 species for metrics combining functional contributions and extinction risk (IUCN rank). **A**-**D**) rankings for different metrics (*see* Methods and Table 1). **E**) Jaccard similarity of the top-10 species across metric pairs. Species’ IUCN status is shown in the lollipops (VU = vulnerable; EN = endangered; CR = critically endangered).

Neither FUS nor FUSE was dominated by a single component used in its calculation: thus, the composite metrics integrate unique aspects of the individual functional indices. All of the top-10 FUS species, and 18 out of the top-20 FUS species, were in the FUSq list, i.e., had values of FUn and FSp that were in the top 10% of values (see Fig. S2 for FUSq species). For FUSE, nine of the top-10 and 12 of the top-20 species were in the FUSEq list (Dataset S1).

### Effects of high-scoring species on functional richness across metrics

Random removal of 10 or 20 species led to 2 and 4% of functional richness loss, respectively (Fig. 4). Removal of the top-10 FUn, FSp and Fdist species resulted in a 19, 33 and 28% loss of functional richness, respectively. Removal of the top-10 FUS species resulted in a loss of 32% of functional richness (Fig. 4). The top-10 FUSq were the same as the top-10 FUS species (Fig. S2) and therefore reduced functional richness by 32% (Fig. S6). Across metrics, the relative impacts of top-10 species varied when weighted by endangerment: the top-10 FUGE and FUSE species held 19% and 15% of functional richness, whereas the top-10 FSGE and FDGE species held only 5 and 3%. These patterns were maintained – albeit with larger losses of functional richness – when 20 species were removed (Fig. 4).

**Figure 4.**
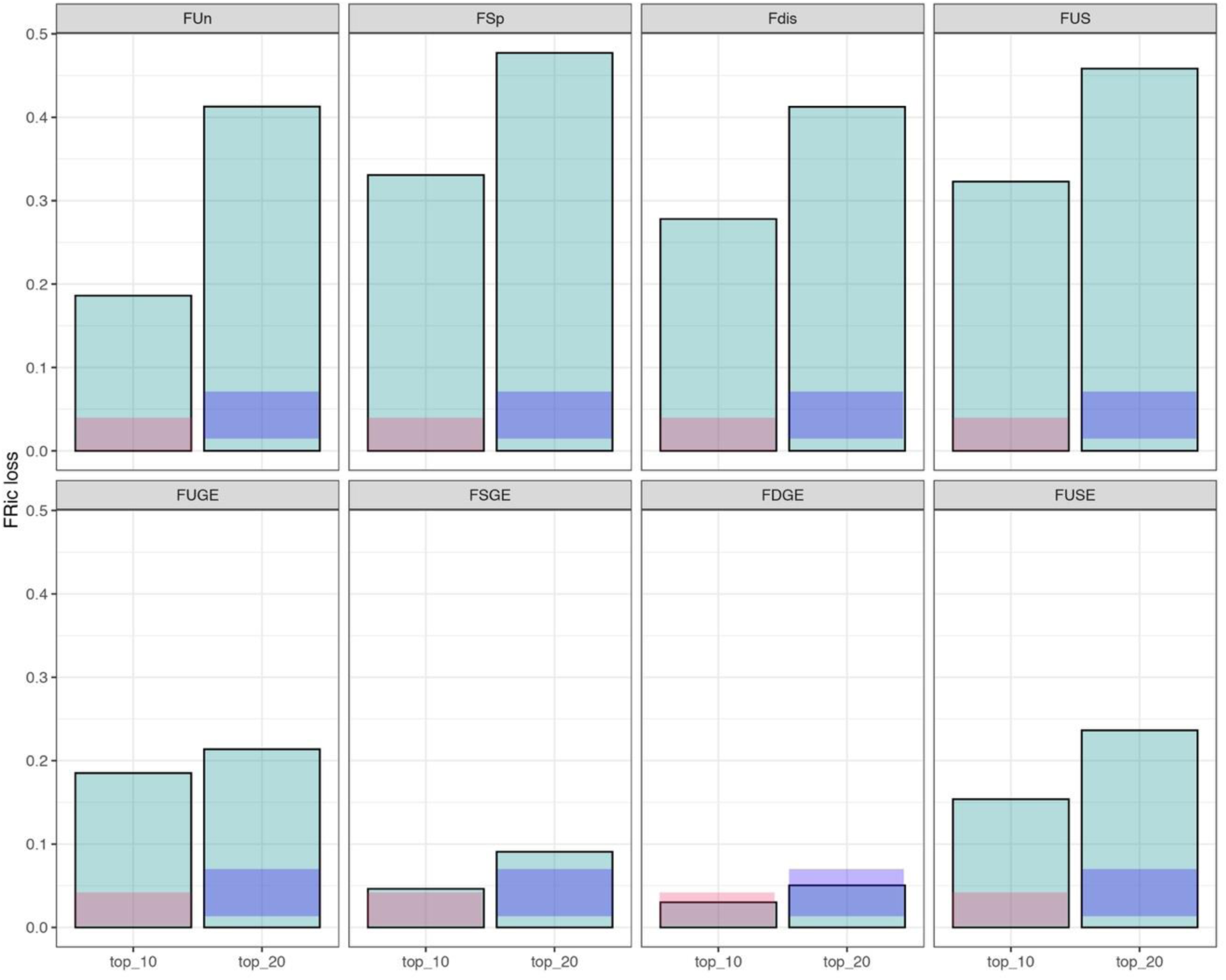
Impact of top-ranking species (top 10 or top 20) for metrics describing functional contributions alone (top panels) and combined with extinction risk (bottom panels) on functional richness. Y-axis is the proportional loss of functional richness. Faded salmon and purple bars show mean +/-1SD of functional richness loss when equal numbers of random species are removed (n = 1,000 iterations). See Table 1 and main text for definition and description of metrics.

### Sensitivity of FUSE to alternative formulations

The alternative formulations of FUSE – whether based on the simple sum of FSGE and FUGE or through the two different direct formulations – were almost indistinguishable (Fig. S7; *r* > 0.99). There was very high overlap between FUSE and both FUSE’ (FUSE_alternative) and FUSE’’ (FUSE_alternative_B), with the same top-seven, the same species contained in the top ten, and nineteen of the top 20 species shared (Dataset S1).

Greater differences between metrics became apparent when weighting by IUCN extinction probability as opposed to rank (Fig. 5). Vulnerable and near threatened species were substantially downweighted. Notwithstanding that the relative ranking of species remained rather consistent (*r* = 0.89-0.91), the influence of extinction probability is clear in the scatterplots and in membership of the top-10 (Fig. S8).

**Figure 5.**
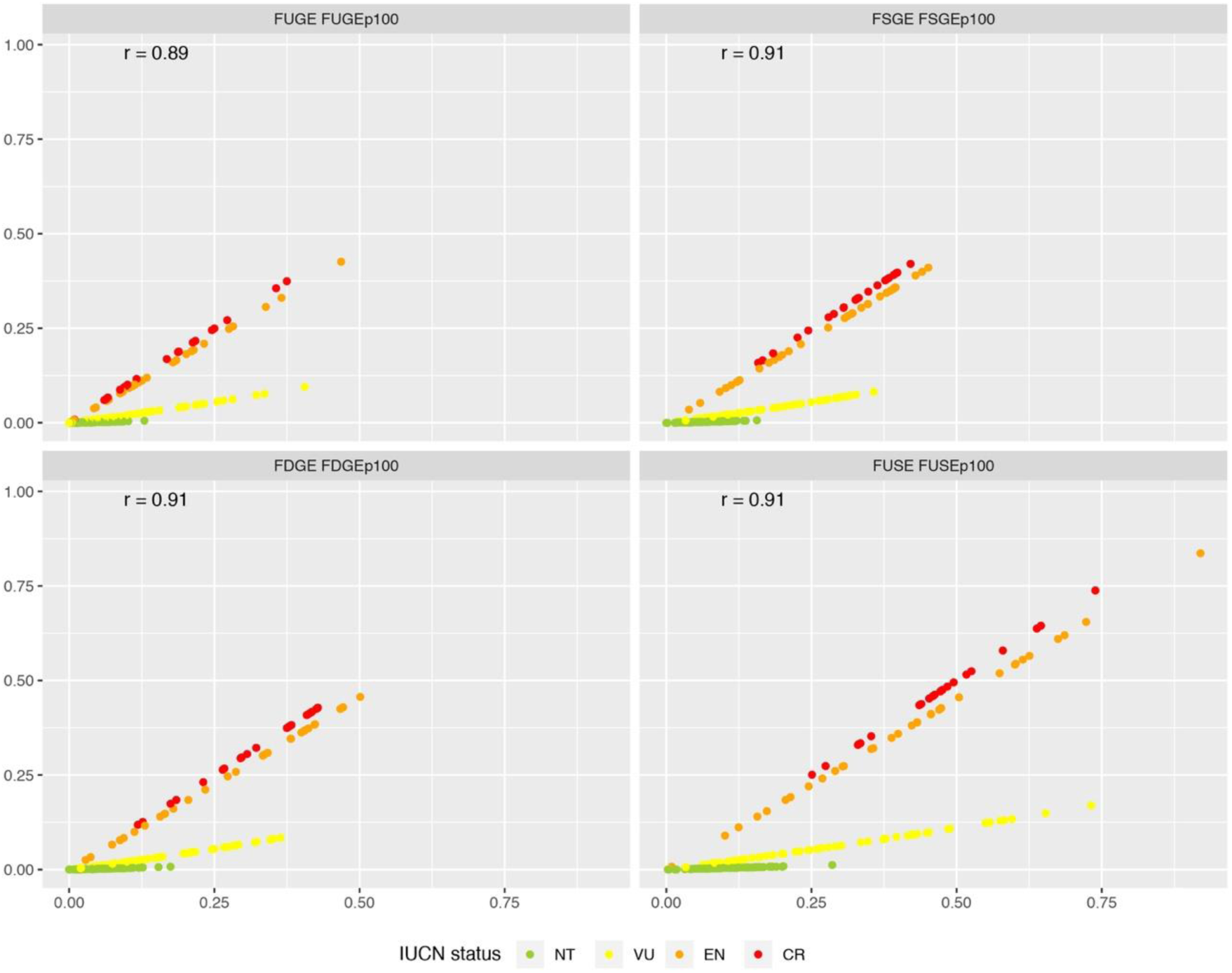
Correlations between metrics weighted with extinction rank (FUGE etc.) and extinction probability (FUGE_P100_ etc.). Colours indicate IUCN status (NT = near threatened; VU = vulnerable; EN = endangered; CR = critically endangered). The Spearman’s rank correlation coefficient is denoted a r. The distinct influence of IUCN status evident in the panels is a result of the different treatment of IUCN status under rank and extinction-weighted metrics. Note that least concern species all score zero for risk-weighted metrics and are not included in this figure.

The impact of removing the top-10 species weighted by IUCN extinction probability (Fig. S8) was less apparent than in the rank-weighted metrics (Fig. S6). This is because the differences in endangerment tend to dominate the species ranking. Consistent with the IUCN-rank weighted metrics, removing the top-10 species for FUGE_P100_ and FUSE_P100_ had slightly stronger effects than FSGE_P100_ and FDGE_P100_ and resulted in losses that tended to be higher than the average of random simulations but not clearly greater than the standard deviation (Fig. S6).

## Discussion

It is now widely recognized that functional diversity is a pivotal biodiversity component to consider when setting conservation strategies (e.g. Devictor et al. 2010; Pollock et al. 2017), especially in the context of the ongoing sixth mass extinction (Ceballos et al. 2015). Our findings confirm that it is important to incorporate multiple metrics of species’ contributions to functional diversity as it changes our understanding of both the total amount of diversity lost as well as the individual species responsible (Mouillot et al. 2013; Violle et al. 2017). Our results provide additional support to our recent publication (Pimiento et al. 2020) and broaden our understanding of the relationships between metrics (Table 1). More particularly, we showed that the FUSE index is robust to different formulations.

Despite a recent emphasis in the literature on functional distinctiveness for setting conservation strategies (e.g. Grenie et al. 2018; Cooke et al. 2020), our results indicate that the complementary information of other metrics should not be overlooked (see also Kosman et al. 2019). Functional uniqueness (and associated IUCN-weighted metrics) captured a different aspect of individual functional contribution vs. functional specialisation or distinctiveness (see also Mouillot et al. 2013; Violle et al. 2017), which appeared to be redundant (see the strong correlation between functional specialisation and functional distinctiveness, Fig. 2). However, these two aspects of functional space are not constrained to co-vary: a species can be functionally distinct to all other species on average but with several neighbours (see *species b* in Fig. 1), implying that its functional role can be compensated in case of extinction. Thus, there is value in combining the unique insights from functional uniqueness and functional specialization to comprehensively evaluate species’ contributions to functional diversity (Buisson et al. 2013; Mouillot et al. 2013).

### FUSE is robust to alternative formulations

The original formulation of FUSE was intended to be a broad barometer of species’ functional uniqueness, specialisation and endangerment as captured by the simple sum of species’ FSGE and FUGE values (Pimiento et al. 2020). FUSE as originally formulated should not be interpreted in terms of a single equation with specific terms for FSp, FUn and GE. Nevertheless, defining FUSE instead as the product of FUS (i.e. the average of FSp and Fun) or FUn and FSp and endangerment (to be more in line with the other metrics used here) creates minimal changes to the rankings of species.

We also evaluated whether FUS(E) could end up emphasising species with exceptionally high values for just a single element used in its calculation. FUS showed strong correspondence with FUSq indicating that it adequately identifies species with high values of both functional uniqueness and specialisation. Notably, analogous results were found for FUSE – especially when the top-10 species were examined. Nevertheless, it may be useful to explore new ways of combining individual functional metrics to emphasise cases where a combination of multiple high values is observed, as in ecosystem function literature (Byrnes et al. 2014).

### Factoring in extinction risk: ranks versus probabilities

Unlike the alternative formulations of FUSE (i.e., FUSE’ and FUSE’’ see Table 1), the treatment of extinction risk (as provided by IUCN status) had a notable influence on the combined metrics, as already pointed out by Mooers et al. (2008) for EDGE. When using probabilities as commonly done in EDGE, the non-linear increase in extinction risk with IUCN status tends to downweight vulnerable and near threatened species. Accordingly, vulnerable species would need to be outliers in the trait space to compensate for the much lower weighting given by IUCN and reach levels of FUSE comparable to endangered or critically endangered species. Note that we used risk probabilities from Mooers et al. (2008) as a simple demonstration of the sensitivity of our indices to treating IUCN as a non-linearly increasing probability rather than a rank. Given the strong influence of extinction probability in metrics such as EDGE (Mooers et al. 2008) and FUSE (this paper), it is obviously vital that extinction probabilities are estimated as accurately as possible in future studies generating extinction probability-weighted metrics such as FUSE_P_ (Davis et al. 2018; Andermann et al. 2020).

### Functionally unique and specialised species support functional richness

Our simulated species removals show that top-ranking sets of species for individual metrics have strong impacts on functional richness. It is not unexpected that the top-ranking species for FSp have the strongest impact on the *richness* facet of functional diversity considered here: these are the species located towards the very edges of the functional space, supporting its full extent. However, it is notable that removing the top-10 FUS species was almost as impactful on functional richness (32 *vs*. 33% loss), despite averaging of functional specialisation with uniqueness. Integrating the extra information on functional uniqueness may help to identify species that are highly specialised *and* unique but not necessarily at the very periphery of trait space. Indeed, as discussed above, high ranking FUS species are high in both FUn *and* FSp. This confirms that high FUS species contribute strongly to maintaining the functional richness facet of functional diversity as argued previously (Pimiento et al. 2020). As FUS also captures functionally unique species, it also identifies species relatively lacking in redundancy. Therefore, although we did not investigate it here, it is possible that removing high FUS species will also disproportionately affect the other facets of functional diversity (see Boyer and Jetz 2014, Leitao et al. 2016). Overall, this deserves to be studied in the future to better understand how the potential extinctions of the most functionally unique and specialised species affect multiple facets of functional diversity.

Removal of species based on top-rankings in the metrics weighted by IUCN status (e.g., FUSE) did not as strongly affect functional richness. This is to be expected because it ‘dilutes’ the top-ranking of functional metrics (e.g., FUS) by introducing an IUCN weighting that is not necessarily tightly linked to a species’ functional contribution. Nevertheless, there were notable differences in impacts across the metrics which departed from the patterns in the contribution metrics. Top-ranking FUGE and FUSE species had stronger impacts than top-ranking FSGE or FDGE species. These results may reflect a slightly greater extinction risk of functionally unique species (Pimiento et al. 2020) and therefore less compromise between species with high FUn and high extinction risk among the top ranked FUGE and FUSE species.

There is inevitably some loss of information with the averaging approach to generate FUS and ultimately FUSE. It is important to recognise differences in the sets of species highlighted through the individual metrics. For example, the vulnerable *Ursus maritimus* (polar bear) is among the top-10 FUGE species, but is only 26^th^ for FSGE, and so is not in the top-10 for FUSE. We suggest that, given the degree of independence of functional uniqueness and specialisation both in theory (Fig. 1, Mouillot et al. 2013), and in real data (this paper, Cooke et al. 2020), the individual metrics of FUGE and FSGE should be considered alongside information from the more integrative FUSE, perhaps by pooling the top-10 set provided by both as the starting basis for prioritisation.

The risk-weighted metrics (FUSE etc.) successfully identified species that are both functionally important and globally endangered. Taking FUSE as an example, these at-risk species only comprise about 3% of marine megafauna but are all among the top 10% for both FUn and FSp and hold a disproportionate portion (15%) of functional richness. By virtue of being threatened, these species have depleted population sizes and distributions. However, as they are not extinct and many show a strong capacity to recover given abatement of extreme exploitation pressure (Duarte et al. 2020), there is still a chance to restore their populations and unique and specialised roles.

### Proceed with caution

It is essential that values of FUSE and related metrics are interpreted and applied with due caution. First, the choice, assessment and weighting of traits can have important bearings on functional diversity (e.g. Lefcheck et al. 2015; Zhu et al. 2017), as can the quality of the functional space (Maire et al. 2015). Our analysis encompasses a wide range of traits, but it is possible to add, refine, and subset traits in any functional diversity analysis according to the ecological question and management priority.

Second, FUSE-like metrics should be considered alongside other relevant information. For example, although the functional roles of many species – including marine megafauna – remain poorly understood, some species have already been identified as playing a keystone role in structuring ecosystems (Estes et al. 2011). Furthermore, established and complementary methods exist for capturing the contribution of endangered species to evolutionary diversity (EDGE, Isaac et al. 2007; Mooers et al. 2008; see also Davis et al. 2018). Future works are clearly needed to evaluate the extent to which the EDGE and FUSE approaches are complementary and can help prioritize the conservation of endangered species.

Third, FUSE-like metrics consider extinction risk and functional contributions simultaneously; however, managers and policy makers may wish to use extinction risk as a first level of prioritization, before examining the relative values of species’ functional diversity contributions *within* each category of IUCN (Cooke et al. 2020). Forth, these metrics must be applied flexibly at the scale of interest, as relatively redundant species at a global scale can be highly unique at a more regional or local scale depending on their distributions (Violle et al. 2017). Finally, ranking schemes should not be interpreted as indicating that some species are expendable. Especially among megafauna, redundancy is likely to be negligible.

## Conclusion

No single metric can capture the functional contribution of a species in nature. However, functional uniqueness is an important facet of an individuals’ functional contribution and can be incorporated with other elements to provide a more general assessment of a species’ contribution. Combination of functional diversity analyses with IUCN status has the potential to help policy makers, regulators, and conservation practitioners set conservation goals in the Anthropocene.

## Acknowledgements

We thank T. Gouhier and P. Pillai for their critical examination of the FUSE index as proposed in Pimiento et al (2020) (personal communication), which inspired us to explore different versions of the FUSE metric.

