## Supplement for "Functionally unique, specialised, and endangered (FUSE) species: towards integrated metrics for the conservation prioritisation toolbox"

Griffin et al.

### Supplemenatry Figures

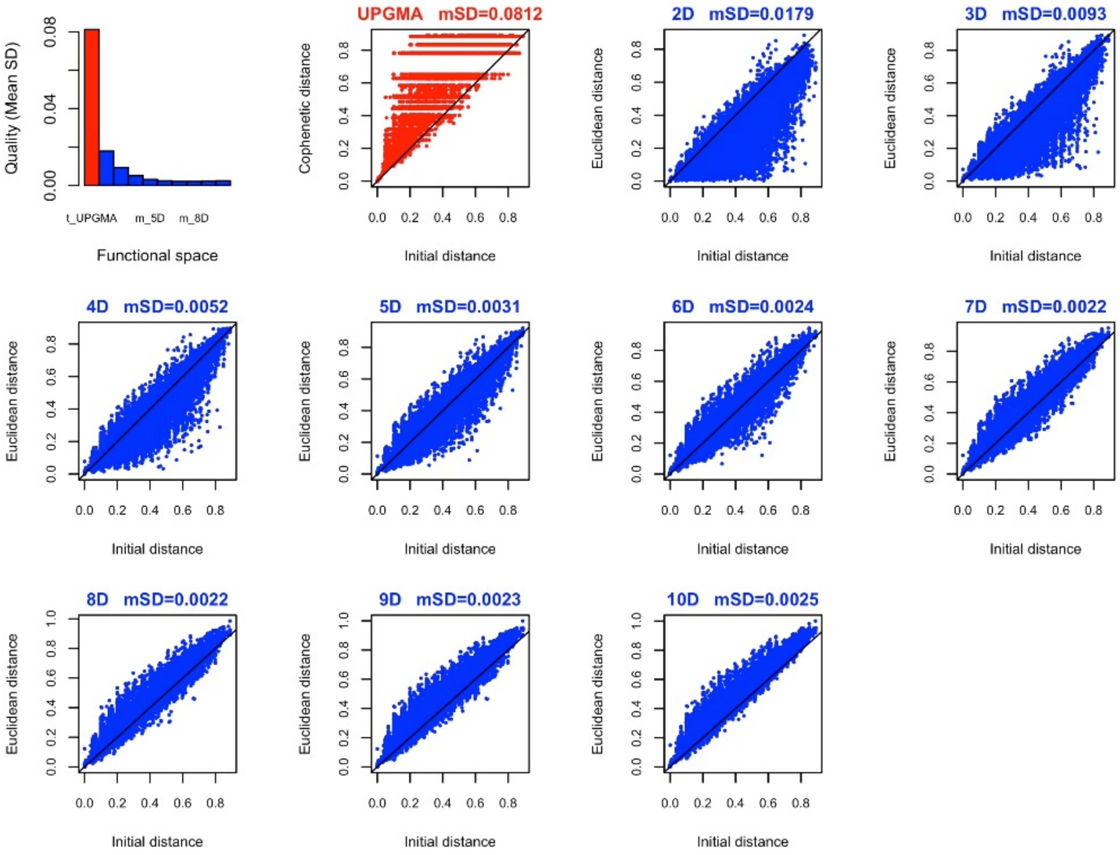


**Figure S1**. Quality of the functional space as captured by the coprrespondence bewtween initital (Gower) distances and the cophenetic distance (for dendrogram approach) and Euclidean distance (for functional space approach used in this paper). mSD is the mean squared deviation between initial trait distance and standardised distance along the deondrogram or in trait space. Following (Maire et al. 2015).

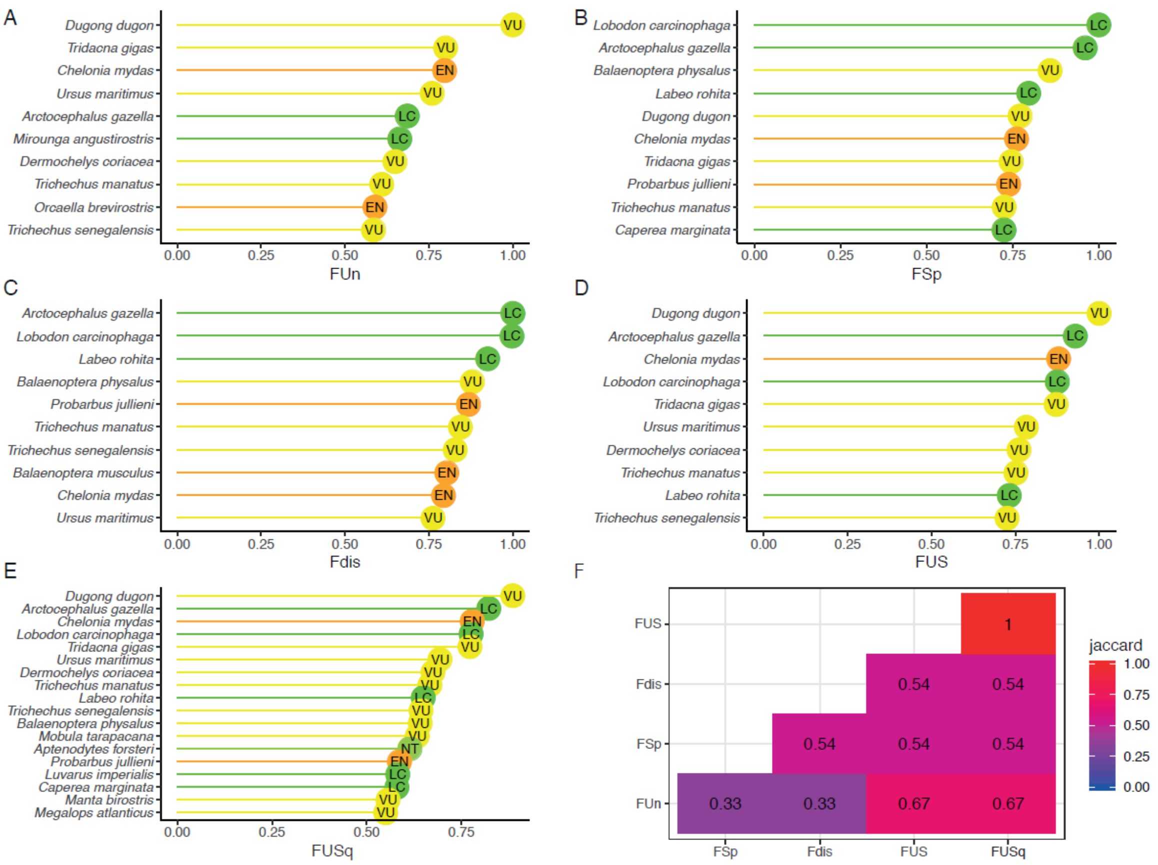


**Figure S2**. Top-10 species for individual functional contribution metrics. Panels A-D show the rankings for different metrics (see Table 1 in main text) and panel E shows the Jaccard similarity of the top-10 species across metric pairs. Species' IUCN status is shown in the lollipops (LC = least concern; NT = near threatened; VU = vulnerable; EN = endangered; CR = critically endangered).

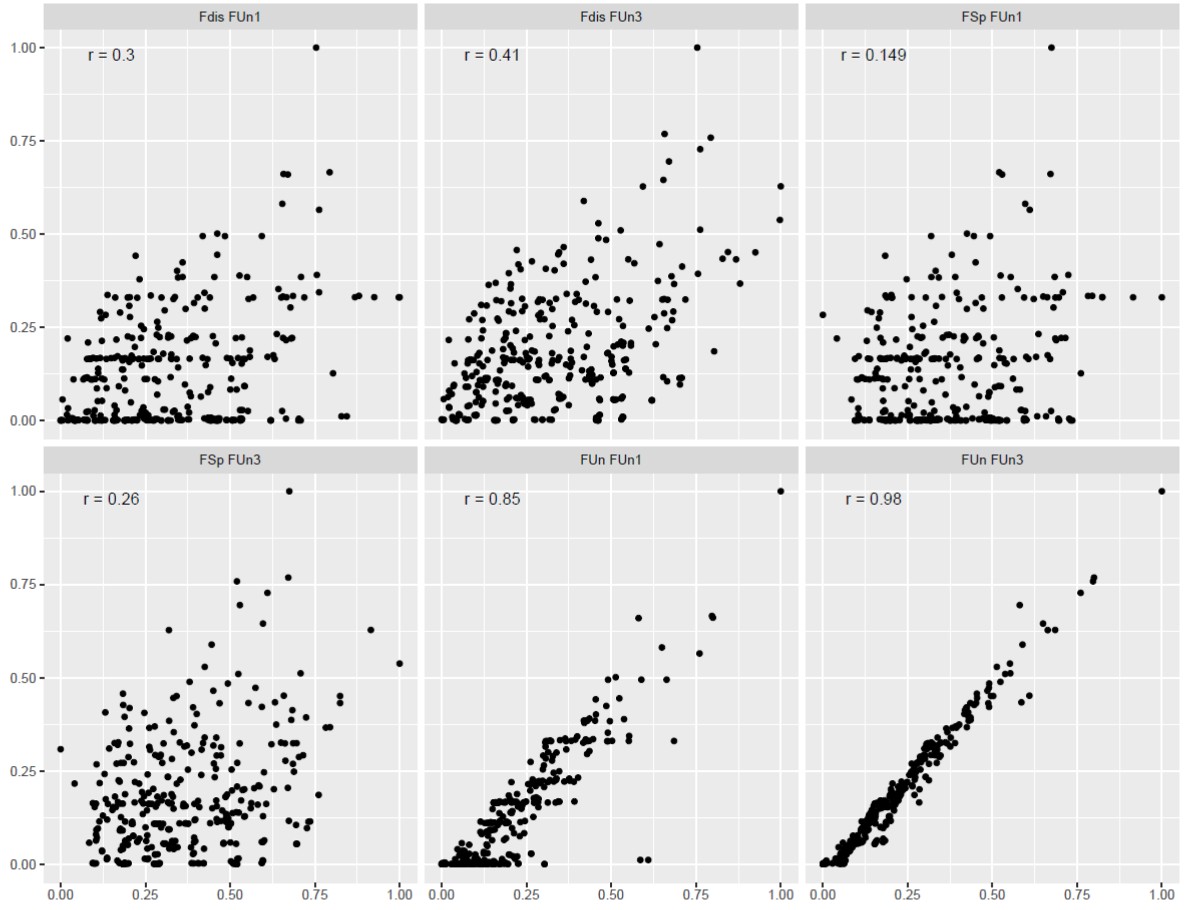


**Figure S3**. Correlations between functional uniqueness (FUn) and other metrics of functional contributions. FUn is functional uniqueness measured as distance to nearest five neighbours, as in the main text and analyses; FUn1 and FUn3 are functional uniqueness measured to the nearest neighbour and three neighbours, respectively. The Spearman's rank correlation coefficient is denoted a r.

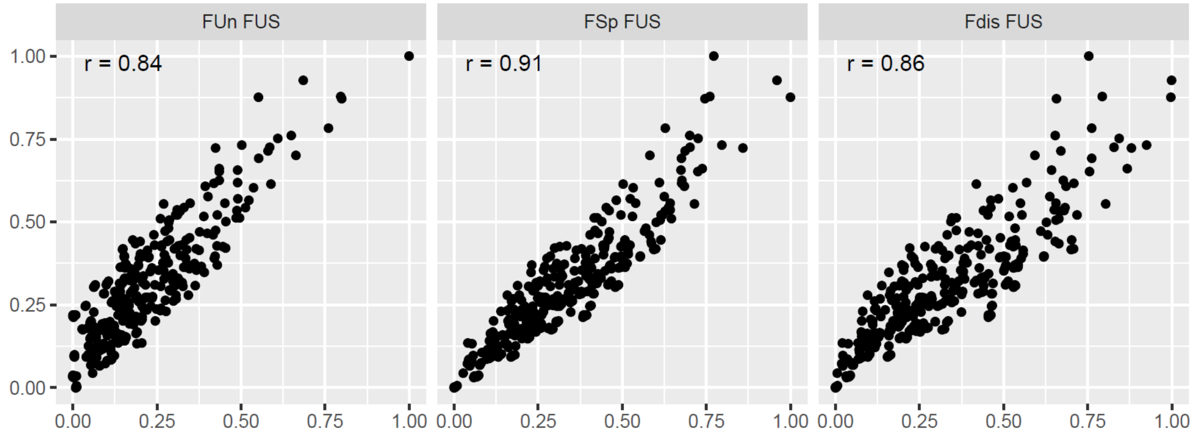


**Figure S4**. Correlations between functional contributions metrics of Fun (functional uniqueness) and FSp (functional specialisation) and functional distictiveness (Fdist) with FUS (the average of functional uniqness and specialisation). The Spearman's rank correlation coefficient is denoted a r.

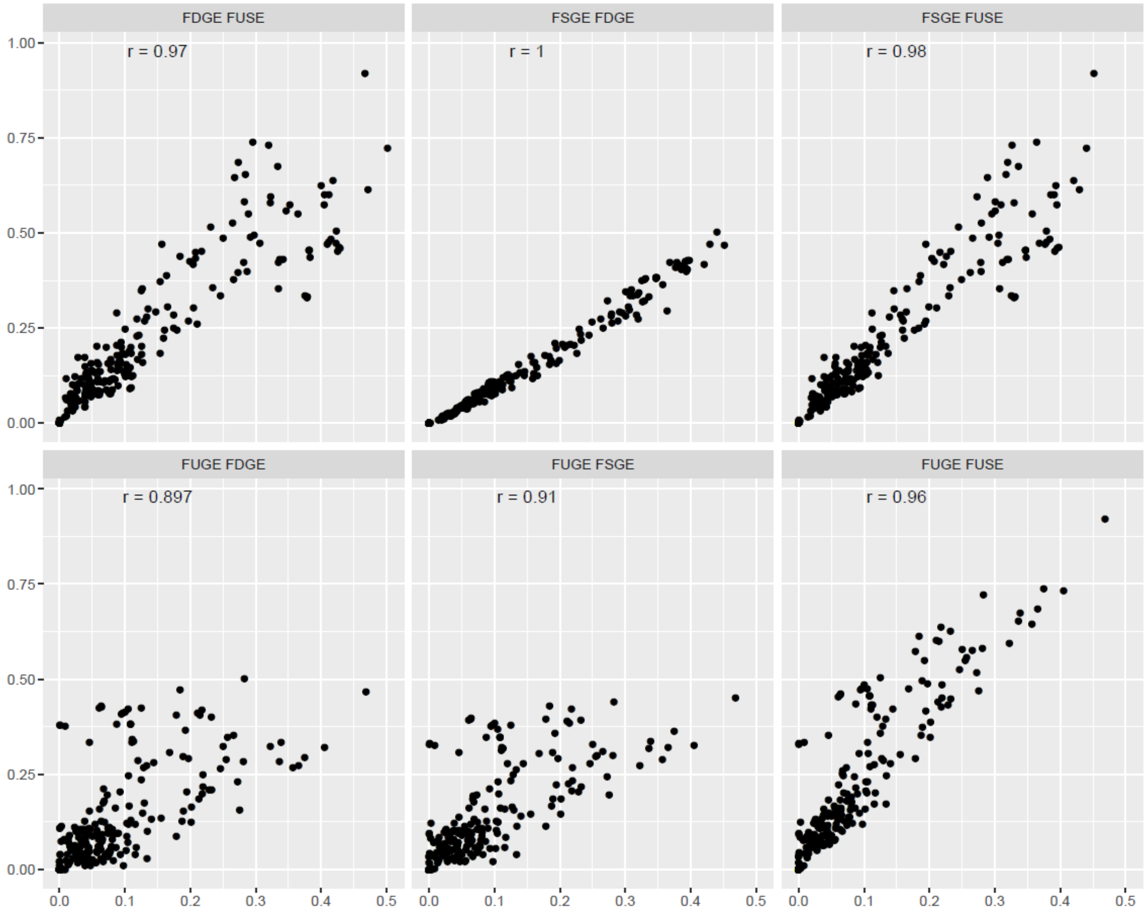


**Figure S5**. Correlations between metrics weighted by IUCN status (rank)
The Spearman’s rank correlation coefficient is denoted a r.

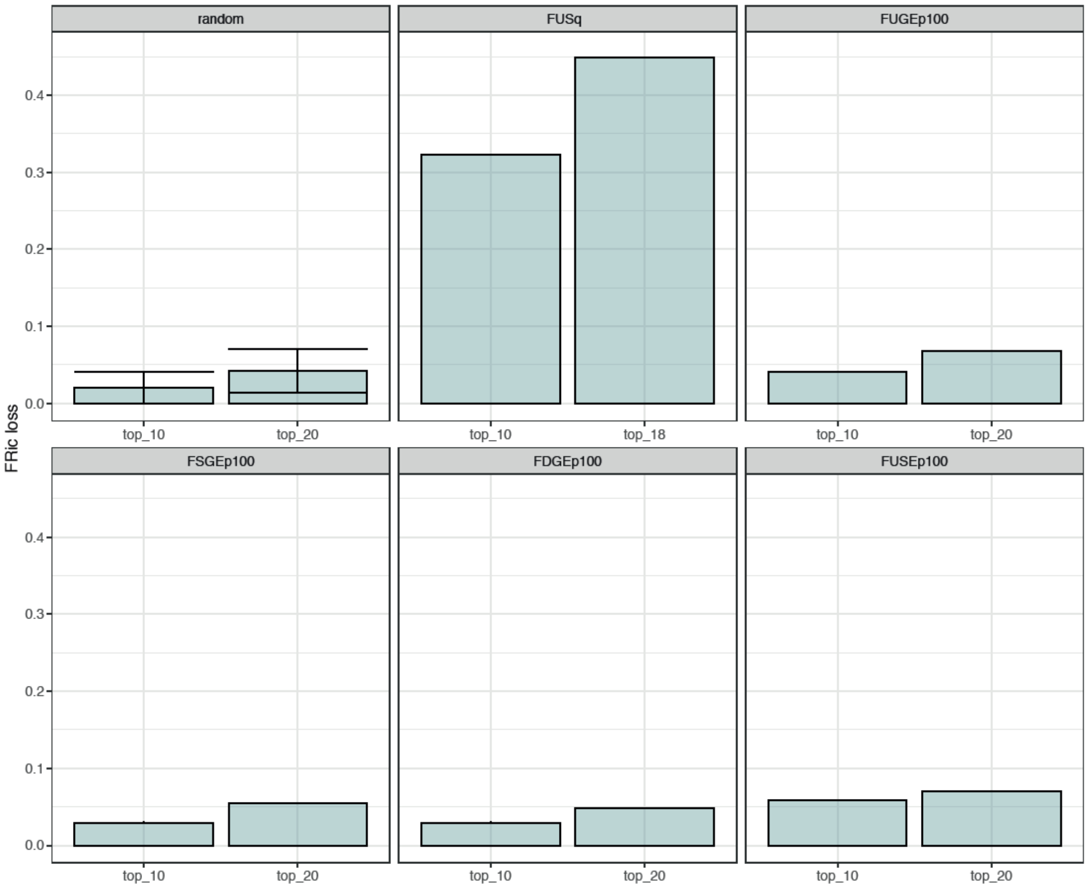


**Figure S6**. Supplementary results showing effect of removing random sets of species vs. top-ranked species for various metrics on functional richness (FRic). FUSq is the list of 18 species in the top 10% for both functional uniqueness and specialisation. $FUGE_{P100}$ etc. show the effect of removing top-ranked species based on functional metrics weighted by extinction probabilities (see Table 1 and main text).

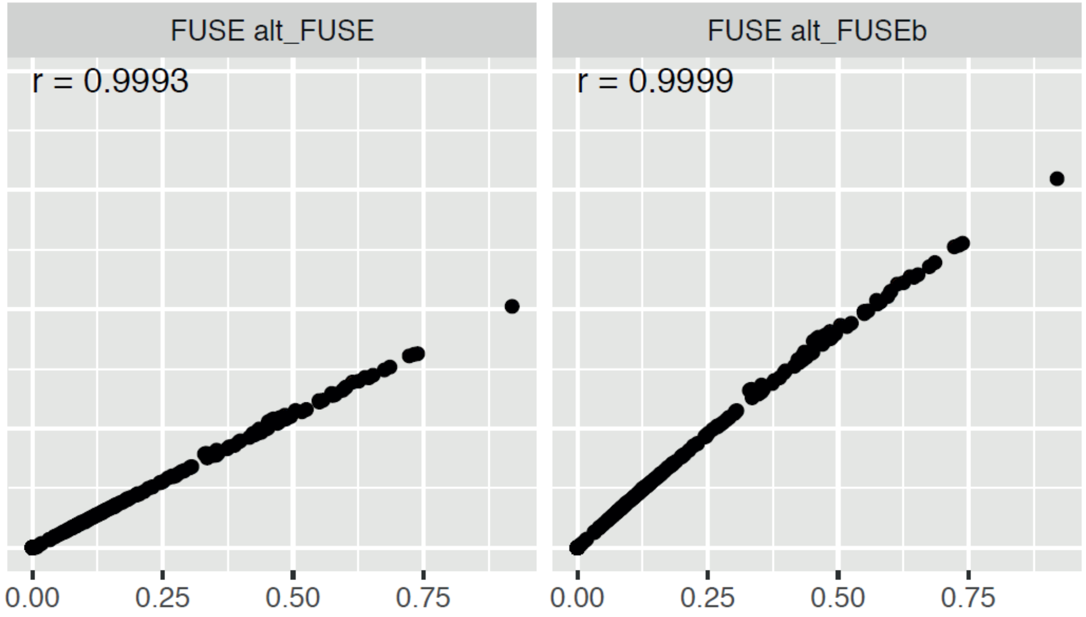


**Figure S7.** Correlations between FUSE as used in (Pimiento et al. 2020) and alternative formulations (see Table 1 in main text). The Spearman's rank correlation coefficient is denoted a r.

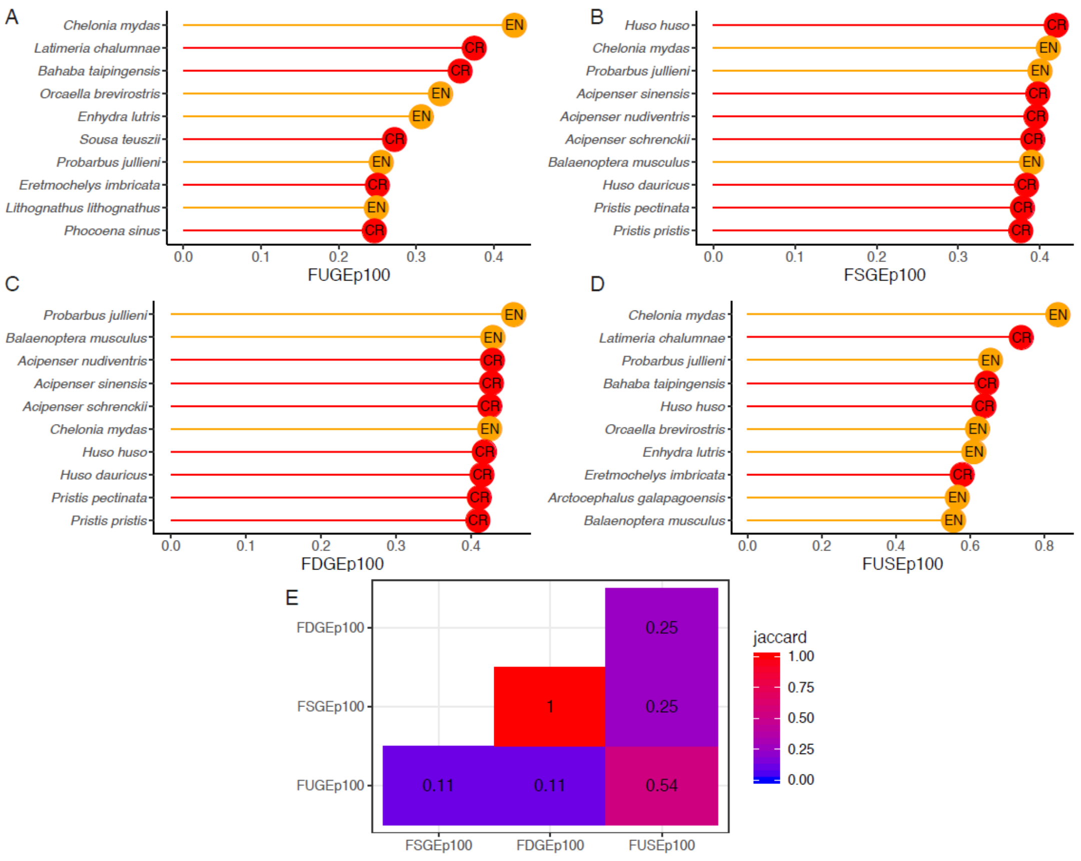


**Figure S8**. Top-10 species for extinction probability-weighted metrics. Panels A-D show the rankings for different metrics (see Table 1 in main text) and panel E shows the Jaccard similarity of the top-10 species across metric pairs. Species' IUCN status is shown in the lollipops (EN = endangered; CR = critically endangered).

**Supplemenatry Datasets**

**Dataset S1.** Marine megafauna species list, their ICUN status and all metrics described in Table 1 (see main text).*Uploaded as a separate file.*

**Cited Literature**

Maire, E., Grenouillet, G., Brosse, S., Villéger, S., 2015. How many dimensions are needed to accurately assess functional diversity? A pragmatic approach for assessing the quality of functional spaces. Global Ecology and Biogeography 24, 728-740.

Pimiento, C., Leprieur, F., Silvestro, D., Lefcheck, J.S., Albouy, C., Rasher, D.B., Davis, M., Svenning, J.C., Griffin, J.N., 2020. Functional diversity of marine megafauna in the Anthropocene. Science Advances 6, eaay7650.
